# TabHLH27 orchestrates root growth and drought tolerance to enhance water use efficiency in wheat

**DOI:** 10.1101/2024.02.29.582695

**Authors:** Dongzhi Wang, Xiuxiu Zhang, Yuan Cao, Aamana Batool, Yongxin Xu, Yunzhou Qiao, Yongpeng Li, Hao Wang, Xuelei Lin, Xiaomin Bie, Xiansheng Zhang, Ruilian Jing, Baodi Dong, Yiping Tong, Wan Teng, Xigang Liu, Jun Xiao

## Abstract

Cultivating high-yield wheat under limited water resources is essential for sustainable agriculture in semiarid regions. Amid water scarcity, plants activate drought response signaling, yet the delicate balance between drought tolerance and development remains unclear. Through genome-wide-association study (GWAS) and transcriptome profiling, we identified a wheat atypical basic helix-loop-helix (bHLH) transcription factor (TF), TabHLH27-A1, as a promising quantitative trait locus (QTL) candidate for both relative root dry weight (DW.R%) and spikelet number per spike (SPS) in wheat. TabHLH27-A1/B1/D1 knockout reduced wheat drought tolerance, yield, and water use efficiency (WUE). *TabHLH27-A1* exhibited rapid induction with PEG treatment, gradually declining over days. It activated stress response genes such as *TaCBL8-B1* and *TaCPI2-A1* while inhibiting root growth genes like *TaSH15-B1* and *TaWRKY70-B1* under short-term PEG stimulus. The distinct transcriptional regulation of TabHLH27-A1 involved diverse interacting factors such as TaABI3-D1 and TabZIP62-D1. Natural variations of *TabHLH27-A1* influences its transcriptional responses to drought stress, with *TabHLH27-A1^Hap-II^* associated with stronger drought tolerance, larger root system, more spikelets, and higher WUE in wheat. Significantly, the elite *TabHLH27-A1^Hap-II^* was selected during the breeding process in China, and introgression of *TabHLH27-A1^Hap-II^* allele improves drought tolerance and grain yield, especially under water-limited conditions. Our study highlights TabHLH27-A1’s role in balancing root growth and drought tolerance, providing a genetic manipulation locus for enhancing WUE in wheat.

## INTRODUCTION

Drought significantly impacts global crop yield, posing an exacerbated challenge due to climate change and intensified human activities (AghaKouchak *et al*., 2021; Ault, 2020; Cook *et al*., 2018). With an anticipated simultaneous drought affecting up to 60% of the current wheat-growing area by the century’s end (Trnka *et al*., 2019), understanding wheat’s response to moderate drought and minimizing yield loss under water deficit is crucial for future food security. Drought impacts wheat growth, development, and yield potential, with susceptibility varying across genotypes and growth stages (Ali, 2019; Mir *et al*., 2012). Critical stages like germination, seedling emergence, tillering, and flowering are particularly vulnerable (Khadka *et al*., 2020). Reproductive stages, when subjected to drought, directly impair phenological and morphological development, leading to reduced yield. Seedling susceptibility is heightened by low soil moisture during emergence, affecting germination, vigor, biomass, and root length, often resulting in failed germination or premature senescence (Ahmad *et al*., 2015; Kizilgeçi *et al*., 2017). Studies across crops have highlighted the close correspondence between drought tolerance in seedlings and adult plants in field conditions. The most resistant wheat cultivars at seedling also among the highest-yielding genotypes in low-rainfall environments (Sallam *et al*., 2018). A close correlation between seedling dry weight and grain yield has been observed in maize and triticale under field conditions (Grzesiak *et al*., 2012).

Plants respond to stress by acclimating their metabolic and physiological processes, achieving a new state of equilibrium. Prolonged stress induces adaptation, altering plant anatomy, growth, and reproduction strategies (Rivero *et al*., 2022). Moisture stress activates intricate drought response pathways, regulating gene expression and signal transduction cascades. Functional proteins like aquaporins and regulatory factors such as bZIP, AP2/ERF, NAC, MYB, WRKY, DREB are triggered (Hrmova and Hussain, 2021; Yang *et al*., 2021). Persistent water deficit induces morphological, physiological, and biochemical changes, including altered photosynthesis and stomatal development, osmotic adjustment, and antioxidant defense. This phenotypic plasticity enables plants to adapt to adverse conditions, ensuring survival and productivity in stressful environments (Rivero *et al*., 2022; Hrmova and Hussain, 2021; Yang *et al*., 2021).

In contrast, water use efficiency (WUE) differs from drought resistance by prioritizing the balance between maximizing yield and minimizing water consumption, vital for crop production improvement (Leakey *et al*., 2019; Tardieu, 2022). Physio-morphological traits like reduced transpiration (small and waxy leaves, deep and sunken stomata), enhanced water absorption capacity (deep, branched and hairy root), and increased harvest yield (coordinated yield components) are effective strategies for enhancing WUE in crops (Chai *et al*., 2015; Khadka *et al*., 2020; Tardieu, 2022). Roots, responsible for water uptake and drought signal sensing, significantly influence drought resistance, grain yield, and WUE. Efficient root systems, featuring optimal spatial distribution, well-developed lateral roots, and increased root hair density, correlate with enhanced drought resistance and higher yields in arid environments (Lynch, 2013; Wasson *et al*., 2012). For instance, deep-rooted wheat subjected to water limitations exhibited up to 35% increased grain weight and 38% higher yield compared to shallow-rooted wheat (El Hassouni *et al*., 2018). Therefore, emphasizing the root system’s role is crucial for developing climate-resilient crops and achieving more resource-efficient agriculture.

Extensive efforts to unveil genetic mechanisms for drought resistance, WUE, and root development in crops employ diverse methods like quantitative trait loci (QTL) mapping, genome/transcriptome-wide association studies (GWAS, TWAS), mutation screening, and multi-omics profiling (Bhardwaj *et al*., 2021; Yang and Qin, 2023). Notable findings include *DRO1* in rice via QTL mapping, enhancing root formation under drought (Uga *et al*., 2013). *ZmNAC111*, identified in maize through GWAS, improves WUE and reduces water loss (Mao *et al*., 2015). In wheat, QTL and GWAS are key strategies for dissecting drought resistance, leading to the identification of genes like *TaNAC071-A1* (Mao *et al*., 2021), *TaWD40-4B.1* (Tian *et al*., 2023), *TaDTG6* (Mei *et al*., 2022), *TaSNAC8-6A* (Mao *et al*., 2020), and *TaVSR1-B* (Wang *et al*., 2021). Selecting superior alleles of stress-tolerance genes enhances crop resilience (Hu and Xiong, 2014). Introducing *DRO1* into a shallow-rooting rice recipient rice cultivar through backcrossing, boosting yield under drought (Uga *et al*., 2013). In wheat, introgression of *TaNAC071-A* or *TaWD40-4B* elite alleles enhanced drought tolerance at the seedling stage, respectively (Mao *et al*., 2021; Tian *et al*., 2023). While genes for drought resistance at the seedling stage are identified, genes for the reproductive stage or WUE have limited breeding applications. Further research is imperative for practical gene cloning and application in crops.

In this study, we identified a transcription factor TabHLH27-A1 through GWAS analysis and transcriptome profiling, based on both relative root dry weight at seedling stage and spikelet number per spike during reproductive growth. Knock-out mutants of *TabHLH27-A1/B1/D1* exhibited reduced wheat drought resistance, grain yield, and WUE. TabHLH27-A1 orchestrates root growth and drought tolerance through interactions with various transcription factors. Notably, introducing the elite allele showcased potential in enhancing drought resistance and grain yield, aligning with its positive selection trends in breeding. Our findings underscore the pivotal role of TabHLH27-A1 in balancing growth and drought tolerance, presenting a genetic manipulation locus for improving WUE in wheat.

## RESULTS

### Identification of *TabHLH27-A1* as candidate for controlling wheat WUE through GWAS

Utilizing a diverse panel of 204 common wheat accessions (**Table S1**), we conducted a Genome-Wide Association Study (GWAS) by genotyping with the Wheat660K SNP array. After stringent filtering, 326,418 high-quality SNPs distributed across 21 chromosomes were retained (**Figure S1a**). The panel was subsequently categorized into three subpopulations (**Figure S1b-d**). Assessing the genetic factors influencing WUE in wheat, we examined multiple traits related to seedling development and yield under well-watered (WW) and water-limited (WL) conditions. Specifically, we evaluated the relative dry weight of the root (DW.R%, calculated as DW.R_WL_/DW.R_WW_ with three repeats, P1, P2, and P3) and spikelet number per spike (SPS) across various environments (six environments, E1-E6). The 204 wheat accessions displayed noteworthy variation in DW.R% and SPS. Notably, a mild positive correlation between SPS and DW.R% was observed (**Figure S1e**).

Moreover, we conducted a GWAS for both the DW.R% and SPS. The analysis employed a mixed linear model (MLM) with corrections for population structure (Q matrix with top three principal components, **Figure S1b**) and kinship (**Figure S1f**). Significant association loci (SAL) were defined as SNP clusters (more than three SNPs with log_10_(*P*-value) ≥ 3.0 in less than 1 Mb distance) present in at least half of the environments. This approach identified 17 SAL associated with DW.R% and 10 SAL with SPS (**Figure 1a,b, Figure S2a-c, Table S2**). Genes located within 1 Mb of the leading SNP were considered as candidates, resulting in 328 candidate genes for DW.R% and 295 for SPS (**Table S3**). Notably, these included wheat orthologs of known genes involved in spike development and flowering (*OsMADS14*, *OsLF*, *OsEPF2*) (Kim *et al*., 2007; Wang *et al*., 2013; Xiong *et al*., 2022), root development (*DLT*, *OsMYB2P-1*) (Dai *et al*., 2012; Li *et al*., 2010), and drought tolerance (*OsMYB48*, *OsGRX8*) (Sharma *et al*., 2013; Xiong *et al*., 2014) (**Table S3**). An intriguing discovery was the overlap between a SAL associated with SPS and DW.R% on chromosome 2A, indicating a shared genetic region (**Figure 1c**).

**Figure 1.**
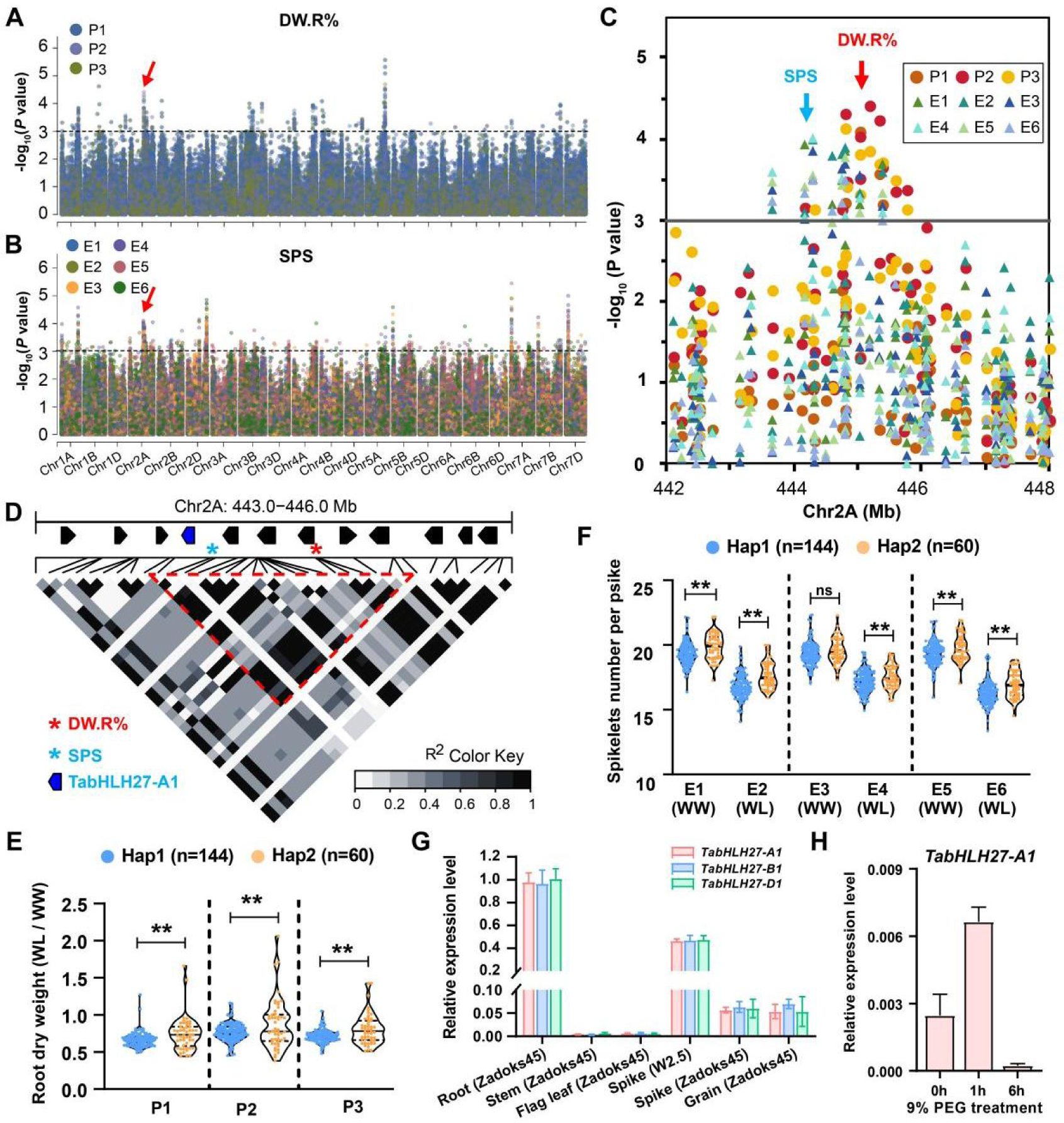
GWAS identified *TabHLH27-A1* as a candidate gene for DW.R% and SPS. **(A,B)** Manhattan plots for DW.R% (**A**) and SPS (**B**) under multiple environments. The y axis refers to −log_10_(*P*). The colors of dots refer to different environments. The loci where *TabHLH27-A1* located was indicated by a red arrow. (C) Local manhattan plot of SNPs in chr2A: 442-448 Mb. The peak SNP for DW.R% and SPS was indicated by a red and cyan arrow, respectively. (D) Heatmap showing linkage disequilibrium (LD) in the 3 Mb physical interval flanking the peak SNPs on chromosome 2A. White to black representing *r^2^* ranging from 0 to 1. The peak SNPs for DW.R% and SPS were indicated with red and cyan asterisks, respectively. The LD block embraced peak SNPs was marked in red dashed triangle frame. *TabHLH27-A1* was marked in blue. **(E,F)** Violin plot indicating the comparison of DW.R% (**E**) and SPS (**F**) among wheat accession with different haplotypes defined by SNPs in the LD block. Wilcoxon rank-sum test was used to determine the statistical significance between two groups. **, *P* ≤ 0.01; ns, no significant difference. **(G)** The spatio-temporal expression analysis of *TabHLH27* in *cv*. KN199 by qRT-PCR with *Tubulin* as the internal control. Error bars show ±SD of three biological replicates. **(H)** *TabHLH27-A1* is induced by short term PEG-mimic drought stress. Two-weeks-old seeding of *cv*. KN199 were treated by 9% PEG (m/v), roots were used for sampling. qRT-PCR were carried out using *Tubulin* as the internal control. Error bars show ±SD of three biological replicates.

Further investigation revealed a 1.67-Mb linkage disequilibrium (LD) block encompassing six high-confidence genes (**Figure 1d**), with SNPs in this block dividing the GWAS panel into two haplotype groups. Accessions carrying Hap2 exhibited significantly higher DW.R% and SPS compared to those with Hap1 (**Figure 1e, f**). Within this LD block, *TraesCS2A02G271700*, a bHLH type TF highly expressed in roots and spikes, exhibits rapid induction followed by decline under osmotic stress, as documented in published transcriptome datasets (Liu *et al*., 2015; IWGSC, 2014) (**Figure S2d-f**) and validated in Kenong199 (KN199) through qRT-PCR (**Figure 1g, h**). Its spatiotemporal expression pattern and responsiveness to drought stress collectively designate *TraesCS2A02G271700* as the most likely candidate WUE gene, subsequently named *TabHLH27-A1*.

### TabHLH27 enhanced the growth and yield of wheat under water-limited condition

The homoeologues of *TabHLH27* exhibit substantial similarity in conserved key protein domains (**Figure S3a**) and share a comparable expression pattern across various tissues (**Figure 1g**). To validate TabHLH27-A1’s role in regulating WUE in wheat, we created two independent mutant lines by simultaneously editing the three homoeologues of *TabHLH27* through CRISPR/Cas9 in wheat *cv.* KN199 (**Figure S3b**). The *Tabhlh27-CR1* line (*CR1*) has premature stops in all three homoeologues, while the *Tabhlh27-CR2* line (*CR2*) has a 24-amino acid deletion in the conserved bHLH domain coding region of *TabHLH27-A1* and premature stops in T*abHLH27-B1* and *TabHLH27-D1* (**Figure S3b**). Both *Tabhlh27-CR1/2* lines displayed slightly inhibited seedling growth under WW conditions and significantly reduced biomass in root and shoot tissues under WL conditions (**Figure 2a,b** and **Figure S4a**). Moreover, *Tabhlh27-CR* lines exhibited heightened sensitivity to drought stress, with a markedly reduced survival rate compared to KN199 after water recovery from severe drought treatment (**Figure 2c,d**). Stomatal density and aperture were not visibly different between KN199 and *Tabhlh27-CR* lines under both WW and WL conditions (**Figure S4b,c**). These findings suggest that TabHLH27 contributes significantly to water limit tolerance.

**Figure 2.**
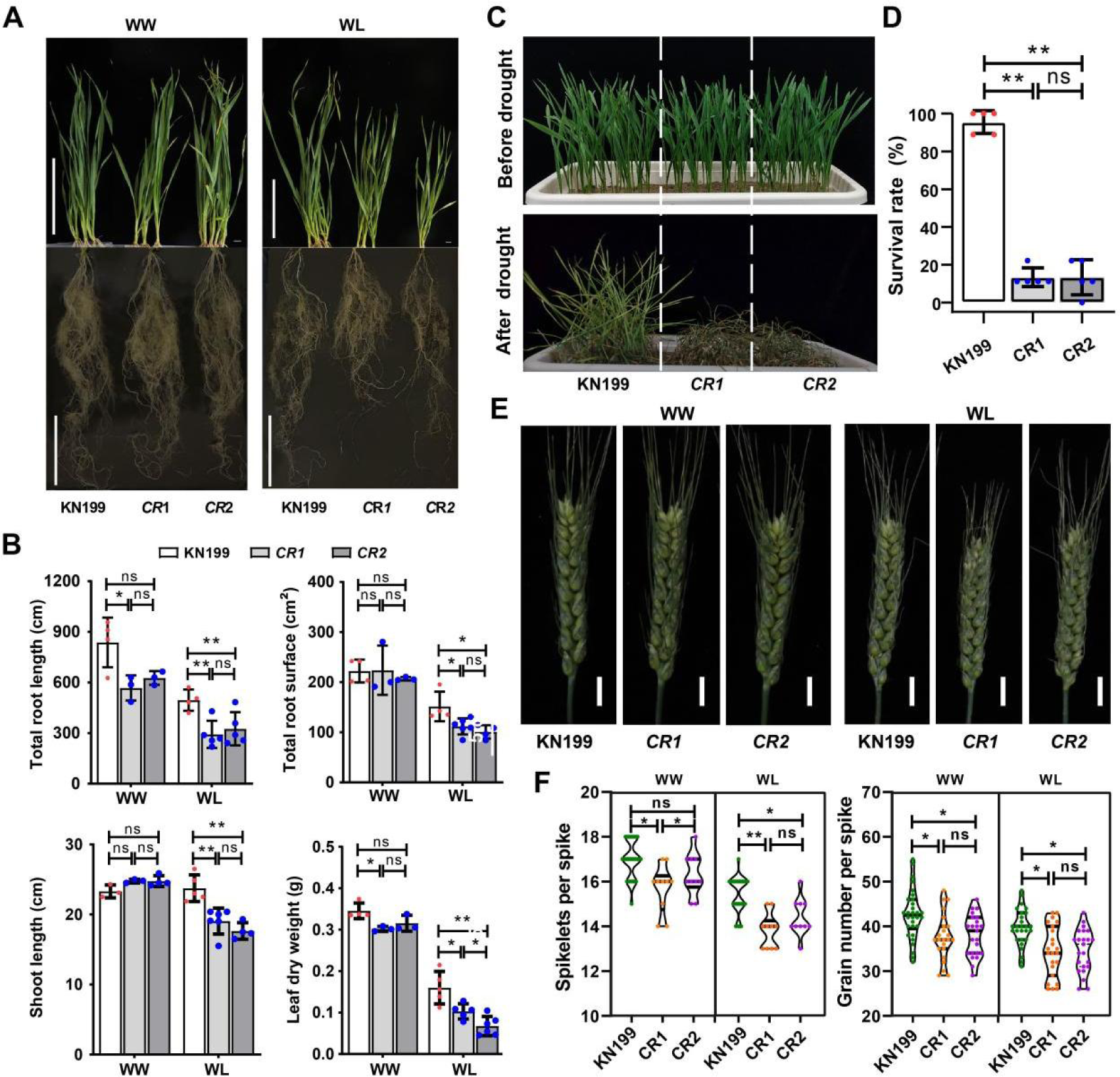
TabHLH27 enhanced the growth and yield of wheat under water-limited condition. **(A)** The shoot and root of *Tabhlh27-CR1, Tabhlh27-CR2* and KN199 under WW and WL conditions. Photos were taken using plants after treatment for one month. Bar=10 cm. (B) Comparison of each trait between *Tabhlh27-CR1, Tabhlh27-CR2* and KN199 under WW and WL conditions. Error bars show ±SD of 4-6 biological replicates. One-way ANOVA (Tukey’s test) were used to determine the statistical significance. *, *P* ≤ 0.05; **, *P* ≤ 0.01; ns, no significant difference. **(C,D)** Assessment of drought tolerance of the *Tabhlh27-CR1, Tabhlh27-CR2* and KN199. Photographs were taken before drought treatment and after a 3-day period of recovery post drought treatment (**C**). One-way ANOVA (Tukey’s test) were used to determine the statistical significance of survival rate between *Tabhlh27-CR* lines and KN199 (**D**). At least 50 seedlings were evaluated for each line, survival rate of five independent replicates were used to compare the significance of the differences. **, *P* ≤ 0.01; ns, no significant difference. **(E)** The spike phenotype of *Tabhlh27-CR1, Tabhlh27-CR2* and KN199 under field WW and WL conditions. Bar = 1 cm. **(F)** Comparison of spikelets per spike and grain number per spike between *Tabhlh27-CR1, Tabhlh27-CR2* and KN199 under field WW and WL conditions. Error bars show ±SD of biological replicates (n ≥ 15). One-way ANOVA (Tukey’s test) were used to determine the statistical significance. *, *P* ≤ 0.05; **, *P* ≤ 0.01; ns, no significant difference.

Given the pivotal role of TabHLH27 in wheat biomass development, particularly during the seedling stage under limited water conditions, we conducted further evaluations of agronomic yield-related traits for *Tabhlh27-a/b/d* mutants in field settings under WW and WL conditions (see methods for detailed WW, WL parameter in field). Under WW conditions, *Tabhlh27-CR2* displayed no distinct phenotypic differences compared to KN199. However, *Tabhlh27-CR1* exhibited a slightly shorter spike length, resulting in fewer spikelets and grains per spike (**Figure 2e,f** and **Figure S4d,e**). Significantly, both *Tabhlh27-CR* lines displayed shortened spikes, fewer spikelets and grains per spike, and reduced grain yield per plant under WL conditions (**Figure 2e,f** and **Figure S4d,e**). Notably, the reduction in these spike-related traits was more pronounced in *CR1* compared to *CR2*, aligning with *CR1* having all three homoeologues mutated whereas *CR2* has a truncated TabHLH27-A1 (**Figure S3b**). For the 1000-grain weight and spike number per plant, there were no significant differences between *Tabhlh27-CR* lines and KN199 under both WW and WL conditions (**Figure S4e**). In summary, these results underscore the role of TabHLH27 in enhancing wheat growth during the seedling stage and increasing grain yield under water-deficit conditions.

### TabHLH27 orchestrates dual function in regulating drought stress response and root development

Under PEG-mimic drought stress, *TabHLH27* showed rapid induction, peaking at 1 hour, and subsequently declined to the pre-PEG treatment levels, becoming nearly undetectable since 72 hours (**Figure S5a**). To investigate its function, we conducted RNA-sequencing (RNA-seq) on root samples from both *Tabhlh27-CR* lines and KN199 after 0, 1, 3 and 72 hours of PEG treatment (**Figure 3a**). Principal component analysis (PCA) revealed a clear separation of samples at 72 hours, while those at 1 and 3 hours were grouped together and distinct from 0 hours (**Figure 3b** and **Figure S5b)**. Upon short-term PEG treatment (within 3 hours), we identified 4,676 differentially expressed genes (DEGs) in KN199, with up-regulated genes (C1, C2, C6, C7) enriched in stress-response Gene Ontology (GO) terms and down-regulated genes (C3, C5, C4) inclined to associate with developmental processes (**Figure S5c,d, Table S4**). Further exploration of PEG-regulated genes with altered expression in *Tabhlh27-CR* lines compared to KN199 revealed a total of 1,077 genes, including 419 up-regulated and 658 down-regulated (**Figure 3c** and **Table S5**). TabHLH27-activated genes (down-regulated in *Tabhlh27-CR*/KN199) were primarily linked to stress-response GO terms, such as “response to water deprivation”, “response to oxidative stress” and “regulation of jasmonic acid mediated signaling pathway”, while TabHLH27-repressed genes (up-regulated in *Tabhlh27-CR*/KN199) were associated with developmental processes (**Figure 3d** and **Table S6**).

**Figure 3.**
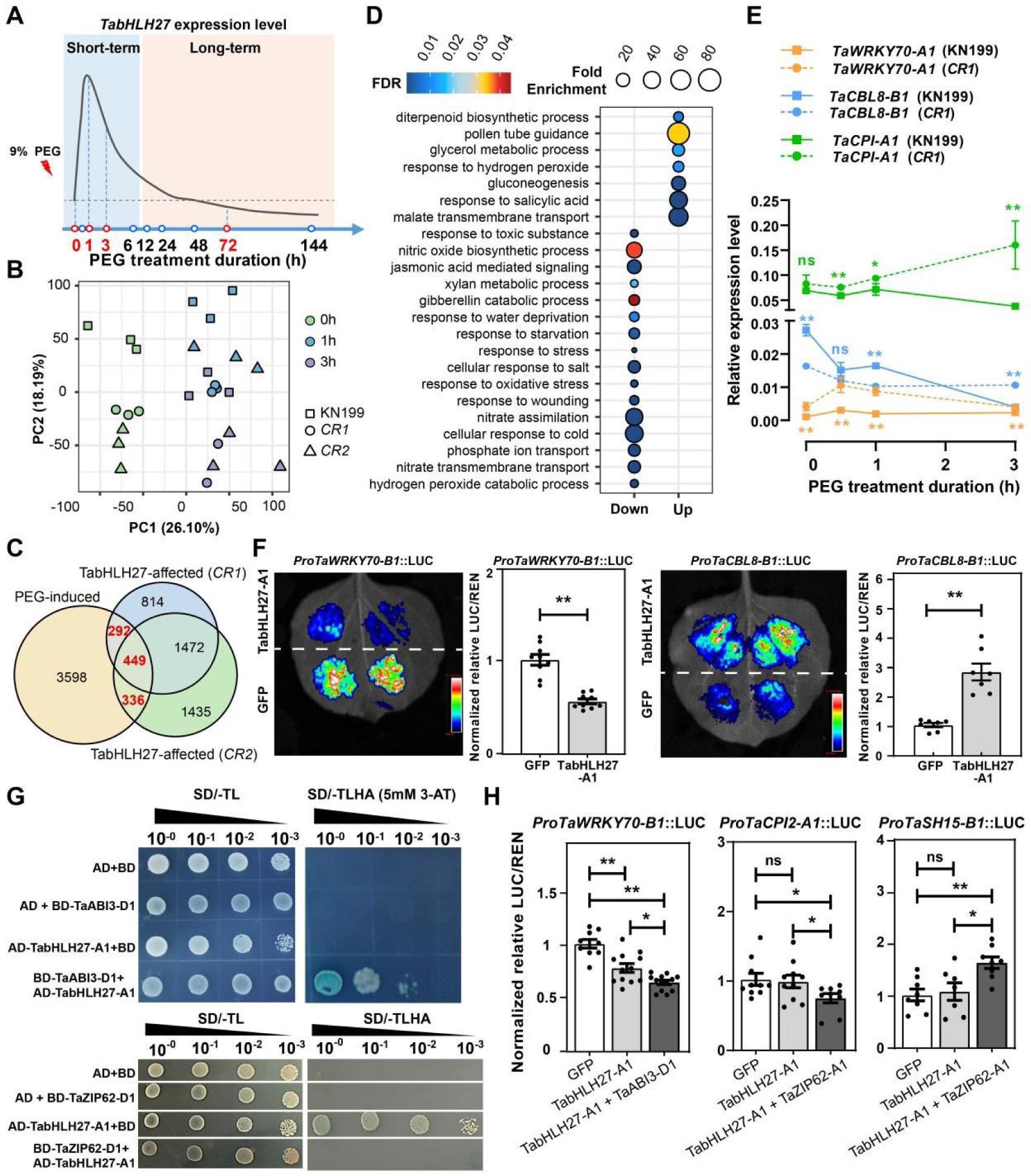
Transcriptome profiling unveils TabHLH27’s dual role in orchestrating both drought stress response and root development. **(A)** A diagram showing experimental design of sampling for RNA-seq. The curved line indicates expression dynamic of *TabHLH27* during PEG treatment for different duration. The dashed horizontal line indicated the basic expression level of *TabHLH27* before PEG treatment. The time points used for RNA-seq sampling were marked in red. h, hour. **(B)** Principal component analysis (PCA) of transcriptome for root samples under short-term PEG treatment. Each sample is represented by a dot, samples from the same time point were in the same color, with KN199, *Tabhlh27-CR1* and *Tabhlh27-CR2* in different symbols. Three biological replicates were sequenced for each line. h, hour. **(C)** Venn-diagram showing the overlapping of PEG-affected genes and TabHLH27-affected genes. The overlapping genes were marked in red. PEG-affected genes indicate DEGs between any two time points in KN199, while TabHLH27-affected genes indicates DEGs between *Tabhlh27-CR* and KN199 in any time point. DEGs weredefined as thresholds of p.adjust < 0.05 and |Fold change| > 1.5. **(D)** Enriched GO terms for TabHLH27-dependent PEG influenced DEGs (genes marked in red in panel c). Fold Enrichment of each term was indicated by the size of dots, with the color indicating the adjust *P*-value. **(E)** The expression levels of *TaWRKY70-B1*, *TaCPI2-A1*, and *TaCBL8-A1* in KN199 and *TabHLH27-CR1* by RT-qPCR. The data are means of at least three independent biological replicates. Student’s t-test was used to determine the statistical significance of expression level at each timepoint between KN199 and *TabHLH27-CR1* (with the color corresponding to the line). *, *P* ≤ 0.05; **, *P* ≤ 0.01; ns, no significant difference. h, hour. **(F)** Dual luciferase reporter assays showing transcriptional regulation of TabHLH27-A1 on *TaWRKY70-B1* and *TaCBL8-B1*. The relative value of LUC/REN was normalized with value in GFP set as 1. Error bars show ±SD of biological replicates (n=7-9). Student’s t-test was used for the statistical significance. **, *P* ≤ 0.01. **(G)** Y2H assay showing the interaction of TabHLH27 with TaABI3-D1 and TabZIP62-A1. **(H)** Dual luciferase reporter assays showing transcriptional regulation of TabHLH27-A1 on *TaWRKY70-B1*, *TaCPI2-A1* and *TaSH15-B1*, when introduced individually or co-transformed with co-factor TaABI3-D1 or TabZIP62-A1. The relative value of LUC/REN was normalized with value in GFP set as 1. Error bars show ±SD of biological replicates (n=7-11). One-way ANOVA (Tukey’s test) were used to determine the statistical significance. *, *P* ≤ 0.05; **, *P* ≤ 0.01; ns, no significant difference.

Interestingly, potential binding motifs of TabHLH27-A1 were found in the promoter chromatin accessible region for wheat orthologs of *WRKY70*, *OsCBL8, OSH15* and *CPI2* (**Figure S5e**). WRKY70 promotes brassinosteroid (BR)-regulated plant growth but inhibits drought tolerance in *Arabidopsis* (Chen *et al*., 2017; Li *et al*., 2013), while OSH15 suppresses panicle size and spikelet number per plant (Wang *et al*., 2022). OsCBL8 and OsCPI2 enhance drought resistance in rice (Gao *et al*., 2022; Huang *et al*., 2007). *TaWRKY70-B1* expression slightly increased following 0.5-hour PEG treatment, significantly up-regulated in *Tabhlh27-CR1* (**Figure 3e**). Luciferase (LUC) reporter assays confirmed transcriptional suppression of *TaWRKY70-B1* by TabHLH27-A1 in tobacco leaves (**Figure 3f**). Conversely, TabHLH27-A1 activated *TaCBL8-B1*, whose expression decreased under PEG treatment and further down-regulated in *Tabhlh27-CR1* (**Figure 3e,f**). Thus, TabHLH27-A1 promotes stress response and inhibits plant growth under rapid drought stress. However, TabHLH27-A1 did not directly regulate *TaCPI2-A1* and *TaSH15-B1* expression, despite the presence of potential TabHLH27-A1 binding motifs in their promoter regions and significant up-and down-regulation in *Tabhlh27-CR* wheat lines (**Figure 3e, Figure S5e,f**).

To unravel TabHLH27’s dual functional transcriptional activity—activating some genes while suppressing others—we probed potential interactions between TabHLH27 and other TFs, forming complexes to regulate downstream genes differently. By scrutinizing coexisting TF binding motifs with TabHLH27 (Min *et al*., 2017; Toledo-Ortiz *et al*., 2003) in the promoter accessible chromatin region (Shi *et al*., 2022) of up-or down-regulated genes, we identified several co-factors with either activation or suppression activity (**Figure S5g)**. Notably, TabZIP62-D1 and TaABI3-D1 emerged as top enriched co-factors for TabHLH27 activated and repressed genes, respectively (**Figure S5h**). The interaction among TaABI3-D1, TabZIP62-D1, and TabHLH27-A1 was confirmed through yeast two-hybridization (Y2H) (**Figure 3g**). Binding motifs of TaABI3-D1 or TabZIP62-D1 were identified in the promoter accessible chromatin region of *TaWRKY70-B1*, *TaCPI2-A1* and *TaSH15-B1* (**Figure S5e**). TabHLH27-A1 individually, suppresses the expression of *TaWRKY70-B1*, with a synergistic enhancement observed when co-transformed with TaABI3-D1 (**Figure 3h**). Co-transforming TabZIP62-D1 and TabHLH27-A1 elicited the suppression of *TaCPI2-A1* expression and the activation of *TaSH15-B1*, while showing no impact when transforming TabHLH27-A1 alone (**Figure 3h)**. Thus, TabHLH27’s diverse transcriptional regulation of drought response and root development genes is likely orchestrated by interacting co-factors.

### TabHLH27 coordinates the short-term stress response and long-term developmental regulation

To gain insights into the dual role of TabHLH27 in coordinating short-term stress response and long-term developmental regulation, we further examined the transcriptome dynamics of root tissue under prolonged PEG treatment (72 hours). We identified 20,735 DEGs between 72 and 0 hours in KN199, primarily enriched in developmental-related biological processes, for instance, “secondary shoot formation” and “regulation of root development” (**Figure S6a,b, Table S7**), indicating the plant’s adaptive growth change during prolonged stress. 1,906 DEGs were found in *Tabhlh27-CR* wheat lines (**Figure S6c, Table S8**), and both TabHLH27-activated and repressed genes were primarily associated with plant growth processes, with stress response-related GO terms also present (**Figure 4a**). For instance, *TaMHZ3-A1*, with an orthologue known to inhibit root development in rice (Ma *et al*., 2018), exhibited suppressed expression initially but recovered and was up-regulated over time (**Figure 4b**). The LUC reporter assay confirmed the significant transcriptional activation of *TaMHZ3-A1* by TabHLH27-A1 (**Figure 4c**), while its expression was notably down-regulated in *Tabhlh27-CR1* (**Figure 4b**). Similarly, TabHLH27-A1 was found to suppress the expression of *TaWRKY74-D1*, the wheat ortholog for *WRKY74* (**Figure S6d,e**), a gene that promotes the tiller number, grain weight, and elongation of primary and adventitious roots under Pi starvation conditions (Dai *et al*., 2016).

**Figure 4.**
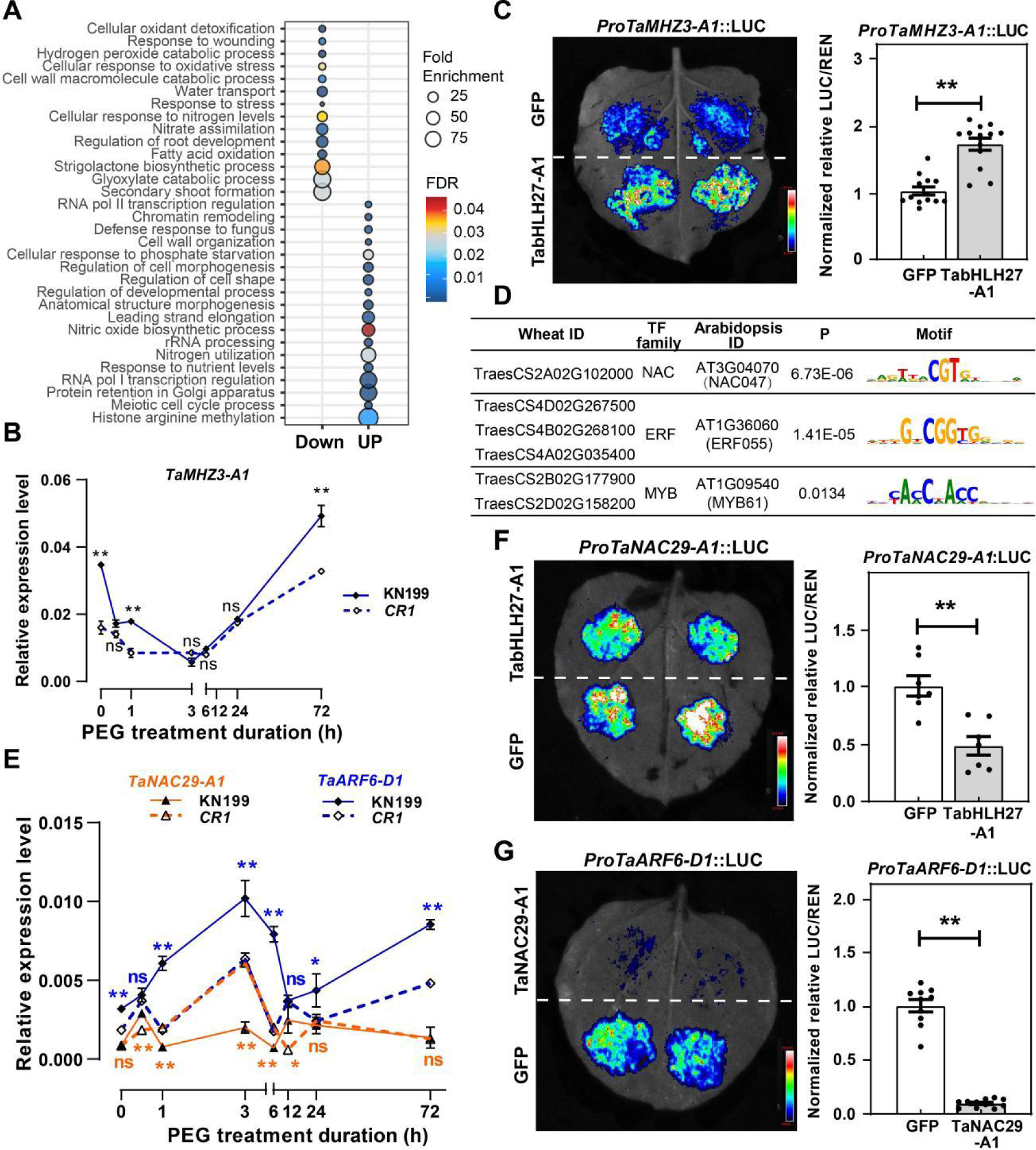
TabHLH27 indirectly regulates root development via mediator under long-term PEG treatment. **(A)** Enriched GO terms for TabHLH27-dependent prolonged PEG influenced DEGs. Fold Enrichment of each term was indicated by the size of dots, with the color indicating the adjust *P*-value. **(B)** The expression levels of *TaMHZ3-A1* in KN199 and *TabHLH27-CR1* by RT-qPCR. The data are means of at least three independent biological replicates. Student’s t-test was used to determine the statistical significance of expression level at each timepoint between KN199 and *TabHLH27-CR1*. **, *P* ≤ 0.01; ns, no significant difference. **(C)** Dual luciferase reporter assays showing transcriptional regulation of TabHLH27-A1 on *TaMHZ3-A1*. Error bars show ±SD of biological replicates (n=13). Student’s t-test was used for the statistical significance. The relative value of LUC/REN was normalized with value in GFP set as 1. **, *P* ≤ 0.01. **(D)** Enriched motifs in the accessible chromatin regions of TabHLH27-dependent PEG influenced DEGs at 72 hours. **(E)** The expression levels of *TaNAC29-A1* and *TaARF6-D1* in KN199 and *TabHLH27-CR1* by RT-qPCR. The data are means of least three independent biological replicates. Student’s t-test was used to determine the statistical significance of expression level at each timepoint between KN199 and *TabHLH27-CR1* (with the color corresponding to the line). *, *P* ≤ 0.05; **, *P* ≤ 0.01; ns, no significant difference. h, hour. **(f,G)** Dual luciferase reporter assays showing transcriptional regulation of TabHLH27-A1 on *TaNAC29-A1* (**F**) and *TaNAC29-A1* on *TaARF6-D1* (**G**). Error bars show ±SD of biological replicates (n=7-11). Student’s t-test was used for the statistical significance. The relative value of LUC/REN was normalized with value in GFP set as 1. **, *P* ≤ 0.01.

Our investigation delved into the hypothesis that short-term targets of TabHLH27, particularly TFs, might contribute to the transcriptional regulation of long-term DEGs identified in *Tabhlh27-CR*. Analysis of motifs enriched in accessible chromatin regions of DEGs at 72 hours revealed enriched NAC, ERF, and MYB binding motifs (**Figure 4d, Figure S6f**), with TabHLH27’s motif ranking less prominently. This suggests the potential involvement of other TFs, like NAC, in mediating *Tabhlh27-CR* induced DEGs at 72 hours. *TaNAC29-A1*, encoding a NAC TF member, was notably up-regulated in short-term PEG treatment in *Tabhlh27-CR* and highly expressed during prolonged PEG treatment (**Figure 4e**). A LUC reporter assay confirmed the transcriptional regulatory circuit between TabHLH27-A1 and *TaNAC29-A1* (**Figure 4f**). Moreover, several DEGs identified in *Tabhlh27-CR* at 72 hours PEG treatment with NAC binding motifs were validated to be activated by TaNAC29-A1 in a reporter assay, including *TaARF6-D1, TaCPI2-A1* (**Figure 4g**, **Figure S6g**). Thus, TabHLH27 emerges as a key regulator in coordinating both short-term drought response and long-term development, utilizing indirect regulation through downstream targets, exemplified by the “mediator” TaNAC29-A1.

### Genetic variations in *TabHLH27-A1* contribute to agronomic traits under water-limit condition

To identify causal variations contributing to superior traits, we sequenced the genomic region of *TabHLH27-A1* (−2Kb of transcription start site to +1Kb downstream of transcription end site) in 32 diverse wheat accessions (Zhang et al., 2017), revealing twelve polymorphic sites, forming two haplotypes (**Figure 5a**). Based on the SNP (−1179) and the 6-bp InDel, the 204 wheat accessions were genotyped and divided into two groups: 153 accessions with *TabHLH27-A1^Hap-I^* (Hap-I) and 51 accessions with *TabHLH27-A1^Hap-II^* (Hap-II) (**Table S9**). Accessions with Hap-II displayed a larger root system and higher transpiration efficiency at the seedling stage (**Figure 5b**). At the reproductive stage, Hap-II produced more spikelets and grains per spike, leading to increased grain yield and WUE compared to Hap-I (**Figure 5c** and **Figure S7a**). The advantages of Hap-II were more prominent under drought conditions. Thus, Hap-II was identified as the superior haplotype with higher drought resistance, higher grain yield and greater WUE.

**Figure 5.**
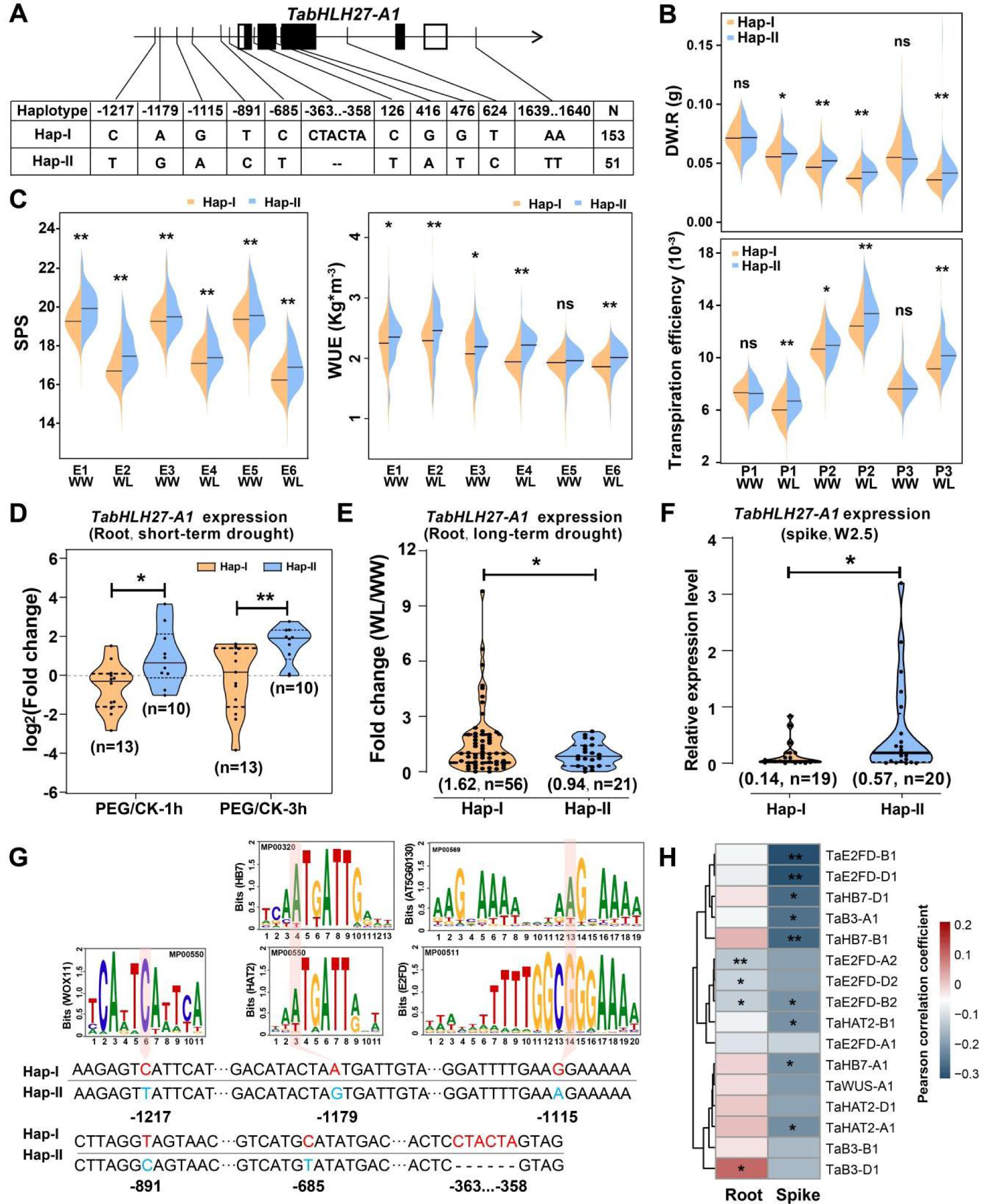
Different haplotypes of *TabHLH27-A1* linked to varied traits and expression level. **(A)** Schematic diagram showing the polymorphism on *TabHLH27-A1* that dividing Hap-I and Hap-II in common wheat population. **(B,C)** Bean plot indicating the comparison of various traits as indicated at the seedling stage (**B**) and reproductive stage (**C**) among wheat accession with different haplotypes defined by SNPs in the genome region of *TabHLH27-A1*. Wilcoxon rank-sum test was used to determine the statistical significance between two groups. *, *P* < 0.05; **, *P* < 0.01; ns, no significant difference. **(D,E)** Violin plot indicating the comparison of expression fold-change between Hap-I and Hap-II. The root of each accession after 1h and 3h PEG treatment was used for RT-PCR (**E**), and roots after growth in soil with water deficit for one month were used for RNA-seq (**F**). Wilcoxon rank-sum test was used to determine the statistical significance between two groups. *, *P* < 0.05; **, *P* < 0.01. The numbers indicate the mean value and sample size for each haplotype. **(F)** Violin plot indicating the comparison of expression level between Hap-I and Hap-II in developing spikes. The spikes at W2.5 stage of each accession were used for RT-PCR. Wilcoxon rank-sum test was used to determine the statistical significance between two groups. *, *P* < 0.05. The numbers indicate the mean value and sample size for each haplotype. **(G)** The predicted TF binding motifs in the promoter sequences of the two haplotypes. The SNPs between two haplotypes were highlighted in the motif. **(H)** Heatmap showing the correlation between *TabHLH27-A1* and potential upstream regulators. The expression level of *TabHLH27-A1* and potential upstream regulators were obtained from RNA-seq data of root samples (n=406, 14 days after germination; Zhao et al. 2023) and developing spikes (n=90, W2.5 stage; Wang et al. 2017). *P* values are determined by the two-sided Pearson correlation coefficient analysis. *, *P* < 0.05; **, *P* < 0.01.

Expression levels of *TabHLH27-A1* were compared between haplotypes. Examining accessions under short-term (via qRT-PCR, 23 cultivars) and long-term (via RNA-seq, 77 cultivars grown for one month with soil moisture restriction) drought conditions revealed Hap-II’s stronger induction with PEG-simulated stress and lower expression during long-term water shortage adaptation (**Figures 5d,e**). Furthermore, Hap-II has a higher expression level in developing spikes (W2.5, 39 cultivars Lin et al., 2024) (**Figure 5f**), aligning with its superior spike traits. We further delved into the contribution of potential causal variations to *TabHLH27-A1* expression by analyzing predicted TF binding motifs in the promoters of two haplotypes (**Figure 5g**). SNPs at positions −1217, −1179, and −1115 disrupted binding motifs of WUSCHEL (WUS), Homeobox 7 (HB7), HOMEODOMAIN ARABIDOPSIS THALIANA2 (HAT2), E2FD/DEL2 factor (E2FD), and B3 domain-containing TF in a chromatin-accessible peak of root tissue (Shi et al., 2022) (**Figure 5g**). Among these, *TaHB7-A1/B1/D1*, *TaE2FD-B1/D1* and *TaHAT2-D1* were induced under short-term PEG stimulus, while others suppressed by drought (**Figure S7b**). Interestingly, the expression level of *TaE2FD-A1/B1/D1* showed a stronger negatively correlation with *TabHLH27-A1* in root (406 wheat accessions, Zhao *et al.,* 2023), while *TaB3-D1* was positive correlated with *TabHLH27-A1* (**Figure 5h**). Similarly, *TaHB7-A1/B1/D1 TaE2FD-B1/D1*, *TaE2FD-B2*, *TaB3-A1* and *TaHAT2-A1* were negatively correlated with *TabHLH27-A1* expression in developing spikes (W2.5, Wang *et al*., 2017). These findings suggest that natural variations of *TabHLH27-A1* affects its transcriptional responses to drought stress, and are associated with drought tolerance, root architecture, spikelets development, and grain yield in wheat.

### Introgression of the *TabHLH27-A1^Hap-II^* allele improves drought tolerance and grain yield in wheat

We further evaluate whether the excellent Hap-II of *TabHLH27-A1* has been selected during the breeding process. In the wheat mini-core collection (MCC) of China, fewer accessions carried *TabHLH27-A1^Hap-II^*, with a lower percentage of landrances possessing the *TabHLH27-A1^Hap-II^* allele compared to introduced cultivars and modern cultivars (**Figure 6a, Table S10**). The frequency of the *TabHLH27-A1^Hap-II^* allele presents a slow-growing trend with cultivar year of release during wheat breeding in China (**Figure S7c**). Hap-I remained the dominant haplotype in most agro-ecological zones of China (**Figure 6a**), except for region VI, indicating that despite being selected during the wheat breeding process in China, the excellent Hap-II of *TabHLH27-A1* has not been widely adopted. Furthermore, the relationship between the proportion of *TabHLH27-A1^Hap-II^* and historical annual rainfall was evaluated using accessions from 14 provinces/cities (the main districts of wheat production) in China. There were fewer accessions carrying *TabHLH27-A1^Hap-II^*in districts with higher annual rainfalls but a higher proportion in districts with lower annual rainfalls, and the proportions of *TabHLH27-A1^Hap-II^* exhibited a strong negative correlation with the annual rainfalls (**Figure 6b**). Therefore, natural variations of *TabHLH27-A1* are linked to enhanced drought tolerance in wheat, the superior *TabHLH27-A1^Hap-II^* was selected during wheat breeding in China, and it still holds potential application in specific areas.

**Figure 6.**
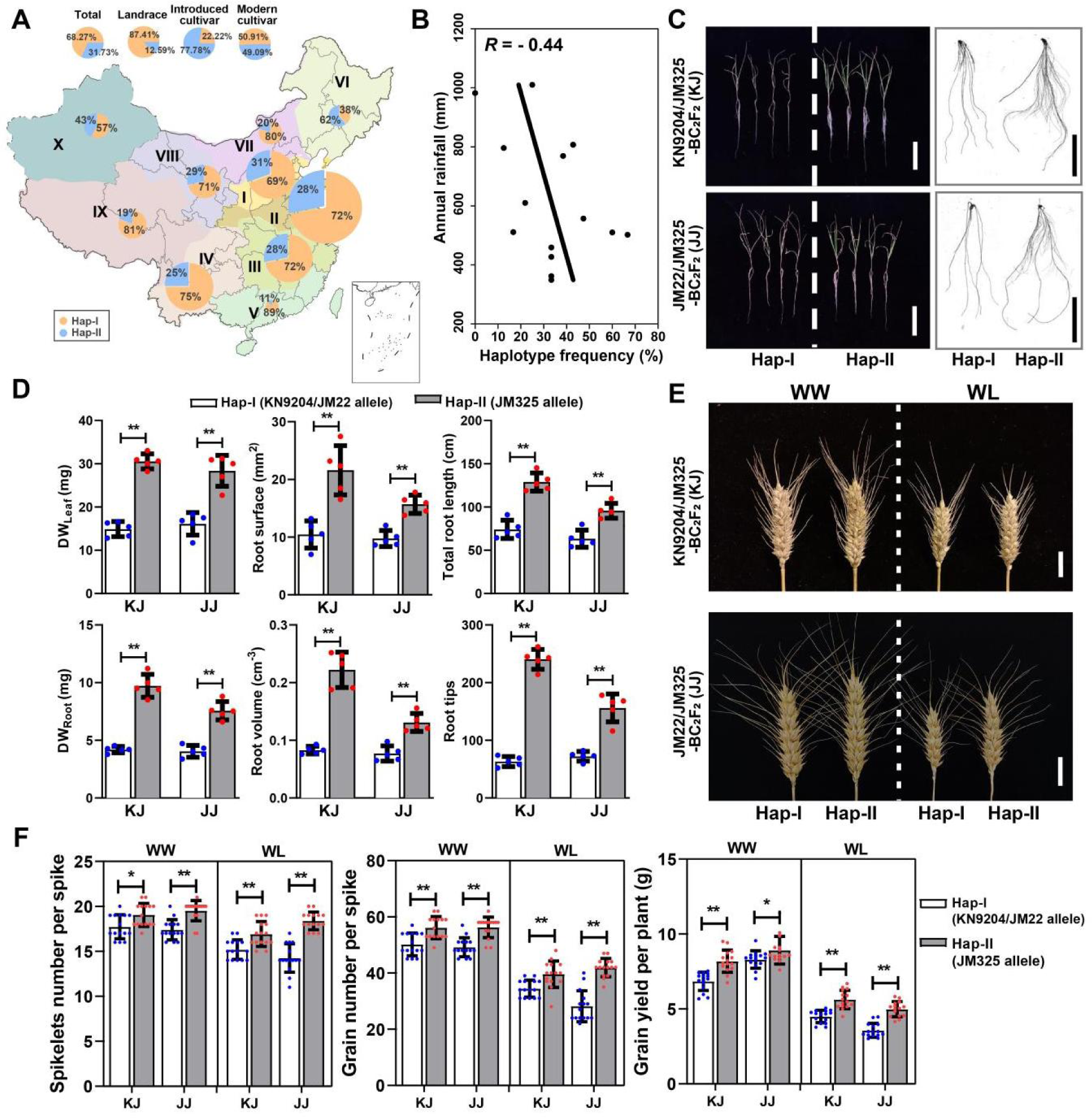
Introgression and assessment of the *TabHLH27-A1* elite allele. **(A)** The percentage of accessions carrying *TabHLH27-A1* Hap-I and Hap-II in different categories and each ecological zones of China. The total accessions were used for haplotype percentage analysis of each ecological zones. The size of pie charts in the geographical map shows the number of accessions, with percentages of the two haplotypes in different colors. **(B)** The proportion of *TabHLH27-A1^Hap-II^* is negatively correlative with annual rainfall with the Pearson correlation analysis (n = 14 wheat planting districts). **(C,D)** The seedling and root system morphology in the BC_2_F_2_ population. After treatment using 9% PEG (m/V) for two weeks, the representative seedlings and roots were taken photos (**B**), and the dry weight of shoot, dry weight of root, and root system architecture were investigated (**C**). Student’s t-test was used for the statistical significance. **, *P* < 0.01. Bar = 10 cm. WW, well-watered; WL, water-limited. **(E)** The spike phenotype of BC_2_F_2_ sibling lines carrying different *TabHLH27-A1* allele under both WW and WL conditions. The sibling lines were planted in greenhouse with water withholding assay. Bar = 1 cm. **(F)** Comparison of SNS, grain number per spike and grain yield between BC_2_F_2_ sibling lines carrying different *TabHLH27-A1* allele under both WW and WL conditions. Error bars show ±SD of biological replicates (n ≥ 10). The student’s t-test was used to determine the statistical significance between two groups. *, *P* ≤ 0.05; **, *P* ≤ 0.01.

To probe whether *TabHLH27-A1^Hap-II^* allele authentically contributes to improving drought tolerance and grain yield in wheat, we conducted a phenotypic comparison between *TabHLH27-A1^Hap-I^* and *TabHLH27-A1^Hap-II^* in wheat, by introducing the *TabHLH27-A1^Hap-II^* allele from the drought tolerant wheat cultivar Jimai 325 (JM325) into two drought-sensitive wheat main cultivated varieties Kenong 9204 (KN9204) and Jimai22 (JM22) carrying the *TabHLH27-A1^Hap-I^*allele. After two times backcrossing of the F_1_ plants, the heterozygous progenies were self-pollinated, resulting in segregating plants in KN9204/JM325 BC_2_F_2_ population (KJ) and JM22/JM325 BC_2_F_2_ population (JJ) carrying either the homozygous tolerant *TabHLH27-A1^Hap-II^* or sensitive *TabHLH27-A1^Hap-I^*allele. Evaluations of drought tolerance using PEG-6000 hydroponics confirmed that the *TabHLH27-A1^Hap-II^* sibling lines have bigger roots and were more tolerant to drought stress at the seedling stage (**Figure 6c,d**). Furthermore, *TabHLH27-A1^Hap-II^* sibling lines of both KJ and JJ populations generated longer spikes with more spikelets, producing more and heavier grains compared to *TabHLH27-A1^Hap-I^* sibling lines under both WW and WL conditions in the greenhouse (**Figure 6e,f** and **Figure S7d**). Collectively, these results illustrate the great promise of the *TabHLH27-A1^Hap-II^* allele for wheat breeding programs with enhancing important agronomic traits under water limitations.

## DISCUSSION

Water limitation profoundly impacts wheat yield, underscoring the need to enhance WUE. Genetic loci governing drought resistance and WUE are pivotal for breeding resilient wheat varieties. While previous efforts predominantly targeted identification factors mediating either seedling or mature stage traits, a comprehensive insight for considering both developmental stages is needed.

### Leveraging joint GWAS analysis for wheat WUE trait understanding

GWAS is crucial for deciphering complex wheat traits like drought resistance (Devate *et al*., 2022; Saini *et al*., 2021). However, current approaches often neglect interconnected traits, hindering comprehensive understanding. Traits evaluating drought resistance, such as root architecture with water acquisition and stomatal conductance with transpiration (Comas *et al*., 2013; Gleason *et al*., 2019), exhibit high correlations. Joint GWAS analysis provides a holistic perspective, integrating genetic data and revealing trait interactions (Gupta *et al*., 2019; Korte and Farlow, 2013; O’Reilly *et al*., 2012). Our study uncovered a positive correlation between DW.R% and SPS (**Figure S1**), identifying a shared QTL (**Figure 1**). Functional analysis of TabHLH27, abundantly expressed in roots and spikes and induced by drought (**Figure 1**), elucidated its role in enhancing drought resistance at the seedling stage and promoting spike development, grain yield, and WUE under water-deficit conditions (**Figure 2**). This exemplifies the potential of joint trait analysis in uncovering multifaceted processes governing wheat’s adaptation to water limitations, crucial for breeding resilient varieties under water-deficient environments.

### Coordinating stress tolerance and plant growth by sophisticated regulation of TabHLH27

In the face of abiotic stresses, plants undergo vital physiological changes, diverting energy from growth to stress defense mechanisms for survival (O’Reilly et al., 2012). However, this trade-off often compromises productivity (Dolferus, 2014). Balancing stress responses with growth is essential, given the energy demands of stress tolerance (Dolferus, 2014; Zhang et al., 2020). Central to this equilibrium are transcription factors, which govern gene regulation by binding to DNA sequences and interacting with other proteins (Pan et al., 2010). Our study on TabHLH27 unveils multifaceted regulation for its dual function in balancing root development and drought stress resistance. Firstly, TabHLH27 exhibits dual transcriptional regulatory activity, activating stress response genes while repressing developmental genes, likely through interaction with different co-factors such as TabZIP62-D1 and TaABI3-D1 (**Figure 3**). Secondly, TabHLH27 shows a dynamic expression profile under drought stress conditions, with rapid induction triggered by PEG treatment, while declining to limited levels under long-term treatment (**Figure 3**). This reduction in stress response may facilitate the lifting of the inhibition of root growth programs for better adaptation to water-limited environments. For instance, down-regulating *TaCPI2-A1*, linked to reduced *TabHLH27* expression, may rely on TabHLH27’s shift from activation to suppression, facilitated by TabZIP62-D1 (**Figure 3**). Thirdly, a circuit regulation between TabHLH27 and other TFs generates a time-course hierarchical transcriptional regulatory network. For example, *TaCPI2-A1*, whose rice orthologs improve drought resistance, is activated by TabHLH27-A1, directly or exemplified by the mediator TaNAC29-A1 (**Figure 4**). This ensures precise regulation across tissues, developmental stages, and dosage levels, as reported in ABA signaling-mediated drought responses in *Arabidopsis* (Song et al., 2016). Therefore, such dynamic regulation highlights the intricate mechanisms of TabHLH27 governing wheat responses to drought stress and root growth stimuli (**Figure 7**).

**Figure 7.**
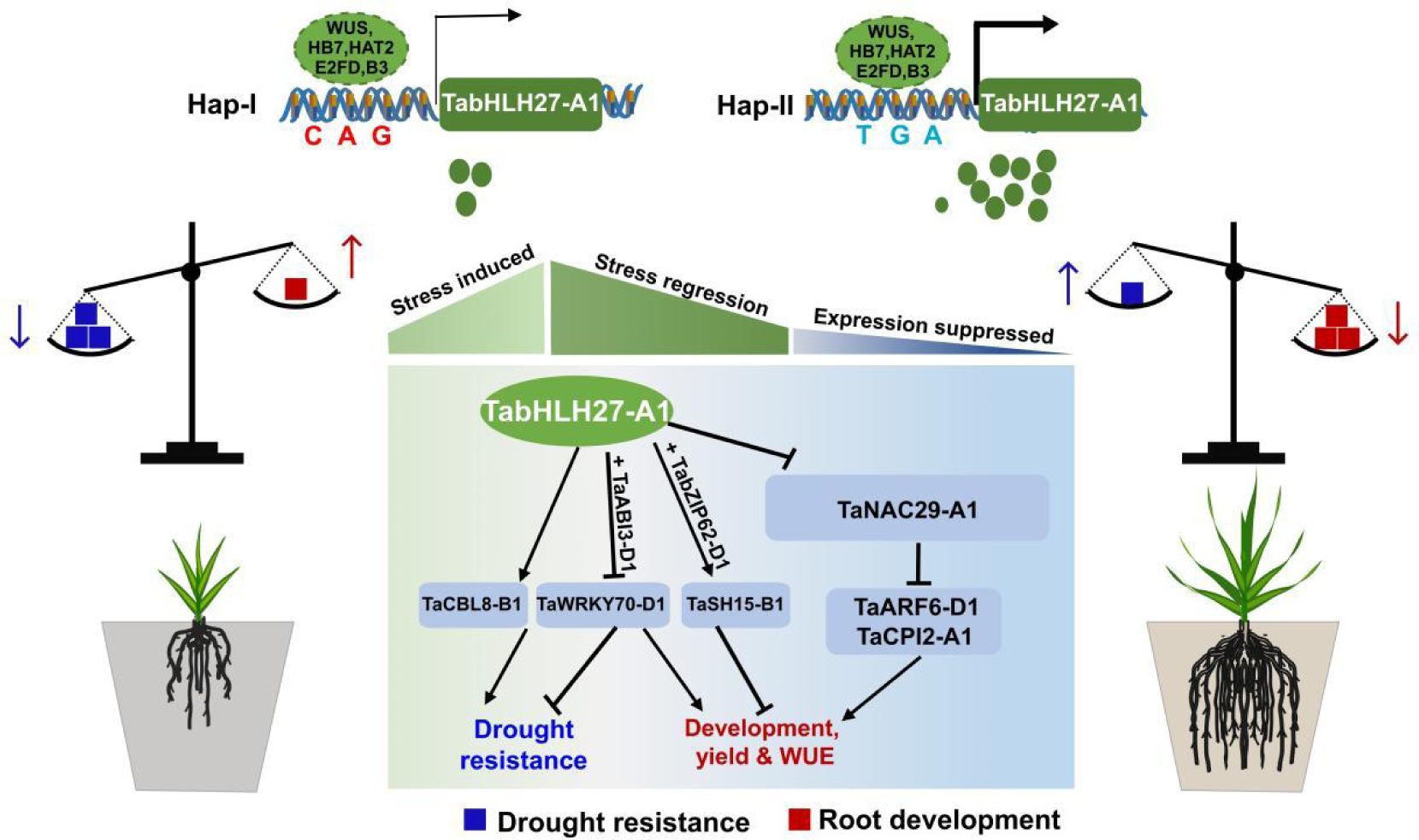
Working model of TabHLH27 mediated drought stress resistance and root growth. TabHLH27 transcriptionally regulate both drought response and root development genes with distinct transcriptional activity, by interacting with diverse co-factors, such as TaABI3-D1, TabZIP62-D1. With the changes of its expression level, TabHLH27 coordinates the short-term stress response and long-term developmental regulation by with the help of “mediator” TaNAC29-A1. Natural variation of *TabHLH27-A1* affect its expression and associated with stronger drought tolerance, a larger root system, more spikelets, and higher grain yield in wheat. Arrow (→) and blunt (┴) in the figure represents activation and inhibition effects, respectively.

### Potential application of elite allele of TabHLH27 for enhancing WUE in wheat

While progress has been made in drought resistance research, particularly in seedling traits, their applicability in breeding remains uncertain. The seedling stage, preferred for its simplicity and cost-effectiveness, is pivotal for studying drought responses and survival rates under extreme conditions. However, genes like *TaNAC071-A1*, *TaDTG6*, *TaSNAC8-6A*, and *TaNAC48* (Chen *et al*., 2021; Mao *et al*., 2020, 2021; Mei *et al*., 2022), primarily linked to survival, lack direct relevance to yield enhancement. Breeding resilient crops necessitates prioritizing stable yield components over survival alone. Striking a balance between both traits is crucial for breeding water-efficient varieties, ensuring moderate productivity while sustaining survival under water stress (Hu and Xiong, 2014; Khadka *et al*., 2020; Rivero *et al*., 2022). TabHLH27 emerges as a key player in balancing drought response and wheat development. The elite allele *TabHLH27^Hap-II^* holds great promise for wheat breeding, enhancing crucial agronomic traits under water limitations (**Figure 5**). Its introgression has demonstrated enhanced tolerance to drought stress at the seedling stage, resulting in increased spikelets and grain production under both well-watered (WW) and water-limited (WL) conditions (**Figure 6 and Figure S7**). The development of a molecular marker for *TabHLH27* allele identification further facilitates molecular-assisted marker selection in breeding practices.

## MATERIALS AND METHODS

### Plant materials and drought resistance evaluation

To knock out *TabHLH27* in wheat *cv.* KN199, two sgRNAs 5’-GCGAACAAGAACATACTGA-3’ and 5’-GTCGTGCCCAACATCACCA-3’ located in the second exon were used. To identify mutations in *TabHLH27-A1* (*TraesCS2A02G271700*), *TabHLH27-B1* (*TraesCS2B02G289900*), and *TabHLH27-D1* (*TraesCS2D02G270300*), genome region around the gRNA targeting sites were cloned using sub-genome gene-specific primers (see **Table S11**) and genotyped by Sanger sequencing.

To access the drought resistance at seedling stage, KN199, *CR1* and *CR2* were subjected to drought stress as described previously with some modification (Mao et al., 2020). The seedlings were grown in greenhouse under 22 °C/18°C (day/night), 16 h/8 h (light/dark), and 40% humidity. Survival rate was recorded after a 3-day period of recovery post drought treatment by scoring all plants with green and viable leaves.

For the observation of plant growth status under moderate drought conditions, one-week old uniform seedlings of KN199, *CR1* and *CR2* were subjected to drought stress following previous reported experimental procedures with some modification (Qiao et al., 2022). A soil moisture content of 15% and 4% was used as a control and drought conditions, respectively. Measurements of stomatal density and aperture were carried out after 3-weeks’ treatment. Epidermal peels from last fully expanded leaf of KN199, *CR1* and *CR2* plants were observed and photographed using an Olympus BX53 microscope. The stomatal apertures were measured and analyzed using ImageJ. Stomatal density defined as number of stomata per mm^2^, and stomatal aperture as width:length ratio of stomatal. The shoot and root of seedling after treatment for one month were harvest separately. Washed roots were scanned by Epson perfection V700 photo instrument and root system architecture indices were obtained using Win RHIZO 2008 software. Then, the dry weight of shoots and roots were measured by drying separately.

Evaluation of drought tolerance under field conditions were carried in Beijing, China (39°54′N, 116°23′E) under field conditions. Each line was planted in three 1.5-m-long rows under WW and WL conditions, with replicates. Plants in WW conditions were irrigated five times throughout the growth period, while the WL conditions stopped watering at jointing stage and exposed to drought stress (with approximately 20-60 % soil water content of WW, degree of drought gradually increase with extension of the treatment). Other agronomic management followed local cultivation practices.

### Expression analysis and RNA-seq

One-week old uniform seedlings of KN199, *CR1* and *CR2* were subjected to PEG-simulated osmotic stress. The roots were harvested after 0, 0.5, 1, 3, 12, 24 and 72 hours treatment in culture solution with or without 9% PEG-6000 (m/v), frozen in liquid nitrogen and stored at −80 °C. Three independent biological replicates were harvested for each sample. Samples from 0, 1, 3 and 72 hours were selected for RNA sequencing.

Total RNA was extracted using HiPure RNA Isolation Kit (Huayueyang, 02160037). First-strand cDNA was synthesized from 2 μg of DNase I-treated total RNA using the FastKing RT kit (TIANGEN, KR116). qRT-PCR was performed using the ChamQ Universal SYBR qPCR Master Mix (Vazyme, Q711-03) by QuantStudio5 (Applied biosystems). The expression of interested genes was normalized to *Actin* gene (*TaActin*, *TraesCS5A02G124300*) for calibration, and the relative expression level is calculated via the 2-ΔΔCT analysis method. Primers used for qRT-PCR are listed in **Table S11**.

### RNA-seq data processing

Raw reads were filtered by fastp v0.20.1 with parameter “--detect_adapter_for_pe, -c, −l 50” for adapters removing, low-quality bases trimming, and reads filtering (Chen *et al*., 2018). Cleaned reads were aligned to IWGSC RefSeq v1.0 by hisat2 with “-5 10 –min-intronlen 20 –max-intronlen 4000” parameters (Kim *et al*., 2019). The raw count of reads of each gene were calculated using FeatureCount software and normalized to TPM (Liao *et al*., 2014). Differential expression genes (DEG) were identified using DESeq2 with a threshold of “p.adjust < 0.05 and |Foldchange| > 1.5” (Love *et al*., 2014). PCA analysis were performed using TPM in FactoMineR (Lê *et al*., 2008). K-means clustering was carried out with “kmeans” function, and heatmap plot using ComplexHeatmap (Gu *et al*., 2016). GO enrichment was performed on http://geneontology.org/ and visualized using ggplot2.

### Luciferase reporter assay

To generate *ProTaWRKY70-B1*::LUC, *ProTaCBL8-B1*::LUC, *ProTaCPI2-A1*::LUC, *ProTaSH15-B1*::LUC, *ProTaMHZ-3-A1*::LUC, *ProTaNAC29-A1*::LUC, *ProTaARF6-D1*::LUC, *ProTaWRKY74*::LUC, we amplified about 2-Kb promoter fragments upstream of each gene from cv. Chinese Spring and ligated them with the CP461-LUC as the reporter vector. The ORFs of *TabHLH27-A1*, *TaABI3-D1*, *TabZIP62-A1*, and TaNAC29-A1 were cloned into the Psuper-GFP vector as effectors, and these plasmids were transformed into *Agrobacterium* GV3101 and injected into *N. benthamiana* leaves in different combinations. Dual luciferase assay reagents (Promega, VPE1910) with the Renilla luciferase gene as an internal control were used for luciferase imaging. The Dual-Luciferase assay reagent (Molecular devices, SpectraMax iD3) was used to quantify fluorescence signals. Relative LUC activity was calculated by the ratio of LUC/REN. The symbol names of the genes and primers used for vector construction are listed in **Table S12** and **Table S11**, respectively.

### Yeast two-hybrid assay

Yeast two-hybrid assays were performed as described in the Frozen-EZ Yeast Transformation II™ (Zymo Research). The coding sequences of *TabHLH27-A1* were cloned into the prey vector (pGADT7), and the coding sequences of *TaABI3-D1*, *TabZIP62-A1* into the bait vector (pGBKT7). The transformed Y2HGold yeast strains were selected on double dropout (Synthetic Dropout Medium/-Tryptophan-Leucine) and quadruple dropout medium (Synthetic Dropout Medium/-Tryptophan-Histone-Leucine). The primers are listed in **Table S11**.

### Identification of the co-factors and mediator of TabHLH27-A1

As the lack of binding motif for TabHLH27-A1, ten closest Arabidopsis orthologs were used alternatively. Protein sequences of all the Arabidopsis bHLH transcription factors with motif information in PlantTFdb were used to construct a neighbor-joining phylogenetic tree, together with of *TabHLH27*.

The bHLH27 up-and down-regulated genes with TabHLH27 motif in the promoter (3 Kb upstream of transcription start site) chromatin open regions (Shi *et al*., 2022), with 50,000 randomly selected sequences in promoter chromatin open regions as background, were used to identify enriched TF motifs (Bailey and Grant, 2021). The TF with enriched TF motifs were considered as co-factors of TabHLH27-A1.

The TF motif enrichment was conducted, by screening motifs in the promoter chromatin open regions of developmental-related DEGs at 72 hours, to identify mediator of TabHLH27-A1. If the enriched TF is a predicted direct downstream target of TabHLH27 (DEG with TabHLH27 motif in promoter chromatin open regions) under short-term drought, it was considered as mediator of TabHLH27-A1 in regulating long-term developmental processes.

### Genotyping, phenotyping and GWAS

The 204 wheat accessions were evaluated at seedling stage and reproductive stage. Cultivars were exposed to WW and WL treatment following reported experiment procedures (Qiao *et al*., 2022), and average root dry weight of each accession was measured. The spikelets per spike was investigated in field at Shijiazhuang (37.85° N, 114.82° E) and Dezhou (37.43° N, 116.35° E) during year 2019-2022. Two replicates were carried out for each accession, in a randomized complete block design. Each block consisted of six three-meters-long rows, with a plant density of 2.7 million ha^-1^. Plots of WW condition was irrigated during the whole life course, and the WL were without irrigation since the jointing stage, and other agronomic management followed local practices. At least 15 representational main spikes in the inner rows were harvest and used for the measurement of spikelets per spike.

Each accession was genotyped using Affymetrix Wheat660K SNP arrays by Capital Bio Corporation (Beijing, China). SNPs were filtered according to the following criteria to get high quality SNP markers: (1) The minor allele frequency is not less than 5%; (2) Missing rate in population does not exceed 10%; (3) Genotype hybrid rate less than 5%; (4) Unique mapped to the reference genome IWGSC RefSeq V1.0. High-quality SNPs of 204 samples were performed to association analysis with phenotypic data, implemented in Tassel v5.2 using the mixed linear model (introducing PCA as a fixed effect and Kship matrix as a random effect in the model). Manhattan plots and quantile-quantile plots were generated using R package “CMplot” (https://github.com/YinLiLin/R-CMplot). Pairwise *r^2^* values were calculated and displayed with LD plots using Haploview 4.2 software (Barrett *et al*., 2005).

### Haplotype analysis of *TabHLH27-A1*

The genomic region of *TabHLH27-A1*, from −2Kb of transcription start site to +1Kb downstream of transcription end site, was sequenced in a diverse set of 32 accessions to identify variations. PCR markers were developed based on the SNP −1179 and the 6-bp InDel, and these polymorphisms were identified for each wheat accession in the natural population to determine the haplotype of *TabHLH27-A1*. The differences of the yield and drought tolerance phenotypes corresponding to different haplotypes were tested.

Natural variation retrieved from the whole-exome sequencing project of the Chinese wheat mini-core collection (Li et al., 2022), were used to assess the breeding selection of *TabHLH27-A1*. The polymorphism with missing rate < 0.5, min allele frequency > 0.05, and heterozygosity < 0.5 were retained for further haplotype analysis. The haplotype frequency in each breeding process of China and among the major Chinese agro-ecological zones were calculated according to the material information provided (Li et al., 2022).

### Introgression of the *TabHLH27-A1* elite allele

Drought-tolerant cultivars Jimai325 (donor parent, carrying the *TabHLH27^Hap-II^*allele) was crossed with drought-sensitive cultivars Jimai22 and Kenong9204 (recurrent parents, carrying the *TabHLH27^Hap-I^* allele), and the obtained F_1_ plants were backcrossed with the recurrent parents for two generations to create the BC_2_F_1_ population. The *TabHLH27-A1* was genotyped in each successive generation, and the heterozygous hybrids were used for backcrossing (genotyping primers see **Table S11**). The heterozygous BC_2_F_1_ plants were self-pollinated, and the resulting BC_2_F_2_ progenies were used for the evaluation of yield potential and drought tolerances.

The drought tolerances of introgression lines at seedling stage were detected by the PEG-simulated stress assay, by culturing one-week old uniform seedlings of BC2F2 population in culture solution with or without 9% PEG6000 (w/v) for 14 days in a growth chamber at 22 °C/18°C (day/night), 16 h/8 h (light/dark), and 50% humidity. To assess the yield potential and drought tolerances at reproductive stage, BC_2_F_2_ plants were planted in greenhouse with water withholding assay. The recurrent parent and its introgression line plants were plant in the same plot under WW condition and WL condition (50% water saving), the plant height, tiller number, spike morphology, and grain yield of representative plants (n ≥ 10) were investigated.

### Statistics and data visualization

If not specified, R (https://cran.r-project.org/;version 4.0.2) was used to compute statistics and generate plots. For two groups’ comparison of data that fit a normal distribution, the student’s t-test was used (Figure 3e, 3f, 4c, 4d, 4f, 4g, 4h, 6c, 6e, S4c, S5a, S5f, S6c, S6d, S6f, S7e). For two groups’ comparison of data that does not fit a normal distribution, the Wilcoxon rank sum test was used (Figure 1e, 1f, 5b, 5c, 5e, 5f, S7a). For three or more independent groups comparison of data, Fisher’s Least Significant Difference (LSD) was used (Figure 2b, 2d, 2f, 3h, S4a, S4e). Pearson correlation was used in Figure 5h and 6b.

### Code and Data availability

The raw sequence data of RNA-seq in this study was deposited in the Genome Sequence Archive (https://bigd.big.ac.cn/gsa) under accession number PRJCA023437. The data analysis method and code are available at github (https://github.com/caoyuan1231/TabHLH27-orchestrates-root-growth-and-droughttolerance-to-enhance-water-use-efficiency-in-wheat).

## ACKNOWLEDGEMENTS

This research is supported by the Strategic Priority Research Program of the Chinese Academy of Sciences (XDA24010204), National Key Research and Development Program of China (2021YFD1201500), Hebei Natural Science Foundation (C2021205013), Full-time introduction of high-end talent research project (2020HBQZYC004), the National Natural Sciences Foundation of China (32100492, U22A6009), Beijing Natural Science Foundation Outstanding Youth Project (JQ23026), and the Major Basic Research Program of Shandong Natural Science Foundation (ZR2019ZD15).

## AUTHOR CONTRIBUTIONS

J.X. and X.-G. L. designed and supervised the research, D.-Z. W. and J.X. wrote the manuscript with the help of X.-X. Z., Y. C. and X.-L. L.; X.-X. Z. did most of the experiments; Y.-P. L., Batool A and X.-M. B. generated the *TabHLH27* knock-out transgenic plants; Y.-Z. Q., B.-D. D., D.-Z. W. and H.W. did all the phenotyping and D.-Z. W. performed GWAS analysis; D.-Z. W. and Y.-P. L. generated the introgression lines and made phenotypic investigation; D.-Z. W. and R.-L. J. did the haploid and selection analysis; Y. C. performed all the bio-informatics analysis with help of Y.-X. X.; D.-Z. W., X.-X. Z., Y.-X. X. and Y. C. prepared all the figures; X.-S. Z., R.-L. J., Y.-P., T., and X.-G. L. revised the manuscript; All authors discussed the results and commented on the manuscript.

## CONFLICTS OF INTEREST

The authors declare no competing interests

## SUPPORTING INFORMATION

**Table S1.** The 204 common wheat accessions in GWAS panel

**Table S2.** Identified QTL for DW.R% and SPS in the GWAS

**Table S3.** List of candidate genes in the QTL region for DW.R% and SPS

**Table S4.** The TPM values of expressed genes

**Table S5.** The DEG list identified in *Tabhlh27-CR* compared to KN199 under short-term PEG treatment

**Table S6.** The enriched GO terms for TabHLH27 activated and repressed genes under short-term PEG treatment

**Table S7.** The enriched GO terms for DEGs in KN199 under prolonged PEG treatment

**Table S8.** The DEG list identified in *Tabhlh27-CR* compared to KN199 under prolonged PEG treatment

**Table S9.** The haplotype of *TabHLH27-A1* in GWAS panel **Table S10.** The haplotype of *TabHLH27-A1* in MCC population **Table S11.** Primers used in this study

**Table S12.** Symbol names of the genes mentioned in this study

## Notes

### Competing Interest Statement

The authors have declared no competing interest.

